# A Bitopic mTORC Inhibitor Reverses Phenotypes in a Tuberous Sclerosis Complex Model

**DOI:** 10.1101/2025.03.14.636465

**Authors:** Sulagna Mukherjee, Matthew J. Wolan, Mary K. Scott, Victoria A. Riley, David M. Feliciano

## Abstract

Neural stem cells (NSCs) of the ventricular-subventricular zone (V-SVZ) generate diverse cell types including striatal glia during the neonatal period. NSC progeny uncouple stem cell-related mRNA transcripts from being translated during differentiation. We previously demonstrated that *Tsc2* inactivation, which occurs in the neurodevelopmental disorder Tuberous Sclerosis Complex (TSC), prevents this from happening. Loss of *Tsc2* causes hyperactivation of the protein kinase mechanistic target of rapamycin complex 1 (mTORC1), altered translation, retention of stemness in striatal glia, and the production of misplaced cytomegalic neurons having hypertrophic dendrite arbors. These phenotypes model characteristics of TSC hamartomas called subependymal giant cell astrocytomas (SEGAs). mTORC1 inhibitors called rapamycin analogs (rapalogs) are currently used to treat TSC and to assess the role of mTORC1 in regulating TSC-related phenotypes. Rapalogs are useful for treating SEGAs. However, they require lifelong application, have untoward side effects, and resistance may occur. They also incompletely inhibit mTORC1 and have limited efficacy. Rapalink-1 is a bitopic inhibitor developed to overcome the limitations of rapalogs by linking rapamycin to a second-generation mTOR ATP competitive inhibitor, MLN0128. Here we explored the effect of Rapalink-1 on a TSC hamartoma model. The model is created by neonatal electroporation of mice having conditional *Tsc2* genes. Prolonged Rapalink-1 treatment could be achieved with 1.5 or 3.0 mg/Kg injected intraperitoneally every five days. Rapalink-1 inhibited the mTORC1 pathway, decreased cell size, reduced neuron dendrite arbors, and reduced hamartoma size. In conclusion, these results demonstrate that cellular phenotypes in a TSC SEGA model are reversed by Rapalink-1 which may be useful to resolve TSC brain hamartomas.

## Introduction

Tuberous Sclerosis Complex (TSC) is a multisystem genetic disorder affecting ∼0.0166 percent of the population^1^. TSC is caused by loss of function mutations in *TSC1* or *TSC2*^2,3^. *TSC1* and *TSC2* encode for the proteins hamartin and tuberin which inhibit Rheb-mTORC1 signaling^4^. *TSC1/2* loss of function mutations activate the mTORC1 pathway^5–8^. mTORC1 is a protein kinase complex that promotes cell growth and proliferation^9^. TSC patients have too much mTORC1 activity and enlarged cells that create tumors called hamartomas within the heart, kidney, lung, skin, and brain^10,11^. Identifying the mechanisms that cause hamartomas to form is important for understanding the many disorders whose genetic mutations affect molecular pathways that intersect with mTORC1 and for developing therapeutic strategies to treat TSC patients.

TSC patients have brain hamartomas called subependymal nodules (SENs) that invade the striatum and subependymal zone (SEZ)^12,13^. SENs are commonly detected during childhood^3,13^. Approximately a quarter of SENs are categorized as subependymal giant cell astrocytomas (SEGAs)^12–14^. SENs may transition into SEGAs^15^. SENs and SEGAs share all histopathological features^16^. Although there is no consensus, the criterion of SEGA diagnosis is >0.5-1.0 cm in size or serial growth^17^. While SEGAs occur throughout the ventricular system, cerebrospinal fluid (CSF) circulation blockade along the caudothalamic groove can cause obstructive hydrocephalus associated with migraines, seizures, and death. Unexplained changes in neurological status or TSC-associated neuropsychiatric disorders (TANDs) can also be a sign of SEGA growth^18^. The median age of SEGA diagnosis is 1 year. Only ∼2.4% of SEGAs are identified after age 40^19^. SEGAs can bleed when being removed leading to non-obstructive hydrocephalus, tissue damage, and mortality^20^. Because SEGAs occur in young children and are located deep within the brain and because surgery comes with a risk, pharmacological intervention is warranted.

mTORC1 inhibitors including rapamycin analogs (rapalogs) are now the standard of care except for cases of acute hydrocephalus^21^. ∼57% of SEGAs are reduced by 50% volume within two years and maintenance doses are not typically associated with changed SEGA volume^18,19,22^. Thus, SEGAs do not always respond to mTORC1 inhibitors. Children may poorly tolerate rapalogs, and if treatments stop, SEGAs grow back^23^. Even surgical removal of SEGAs is followed by regrowth in nearly 40% of patients. The mechanisms that account for SEGA regrowth are unclear but may be related to the fact that mTORC1 allosteric inhibitors incompletely inhibit mTORC1 phosphorylation of select substrates^24,25^. To overcome the limitations of rapalogs, a novel bisteric inhibitor linking MLN0128 to rapamycin called Rapalink-1 was generated^26^. Rapalink-1 simultaneously inhibits mTORC1 through the rapamycin moiety by targeting FK506-binding protein 12 (FKBP12) and the FKBP12 rapamycin binding domain of mTOR as well as inhibiting both mTORCs through the ATP competitive inhibitor moiety, MLN0128^26^. Since mTORC1 controls cell growth and mTORC2 distinctly regulates the actin cytoskeleton, dual inhibition by Rapalink-1 may have advantages over rapalogs.

We created a mouse model of TSC SEGAs by electroporating CRE recombinase into NSCs of mice having conditional *Tsc2* genes to provide mechanistic insight into SEGA pathogenesis^27,28^. Mice developed hamartomas with SEN-like lesions that develop into SEGA-like hamartomas. We previously found that TSC mutant NSC translational programs were altered and prevented differentiation leading to the aberrant production of neurons in the striatum. These lesions were associated with ensembles of cytomegalic neurons and giant cells. We performed single nuclei RNA sequencing (snRNA-Seq) of these mice and discovered altered NSC transitional states caused by loss of *Tsc2*. Moreover, neurons were a core feature in this model and had altered transcriptomes. The extent to which mTOR activity might cause these phenotypes was not assessed.

What follows are the results of a study that utilizes the bisteric inhibitor Rapalink-1 on a TSC model of striatal hamartomas representing SEGA-like lesions.

## Results

### Rapalink-1 Treatment of a TSC Mouse Model

*Tsc2*^*wt/wt*^ and *Tsc2*^*f/f*^ x *RFP* neonatal mice were electroporated with Cre recombinase and GFP encoding DNA plasmids (**Figure 1A**). Electroporation allows for the targeting of lateral V-SVZ NSCs that generate striatal glia (**Figure 1B**). This causes recombination leading to deletion of exons 2-4 of *Tsc2* and red fluorescence along the lateral ventricles that appear as SEGA-like hamartomas in *Tsc2*^*f/f*^ mice (**Figure 1C**). Hamartomas had elevated mTORC1 activity as assessed by pS6 staining (**Figure 1D-I**) as we previously reported^27^. *Tsc2*^*f/f*^ x *RFP* neonatal mice were subsequently randomized, assigned a unique identification number, and treated with Rapalink-1 or DMSO (control) for 30 days or until sacrificed (**Figure 1J**). Animals were euthanized by intraperitoneal injection of euthasol or CO_2_ inhalation followed by swift decapitation. No mice in the control group died and no significant changes in behavior or signs of distress were noted (**Figure 1K**). Cohorts of mice were given 1.5 mg/kg or 3.0 mg/kg Rapalink-1 once every five days. 3.0 mg/kg Raplink-1 treated mice survived on average 88.2 days (N=8). Likewise, 1.5 mg/kg Rapalink-1 was well tolerated, and a single mouse died prior to the scheduled termination date likely because of injection error (N=5) with all remaining mice surviving 90 days. These doses were well tolerated with only the higher dose having slightly lower weights at the end of treatment (**Figure 1L**).

**Figure 1.**
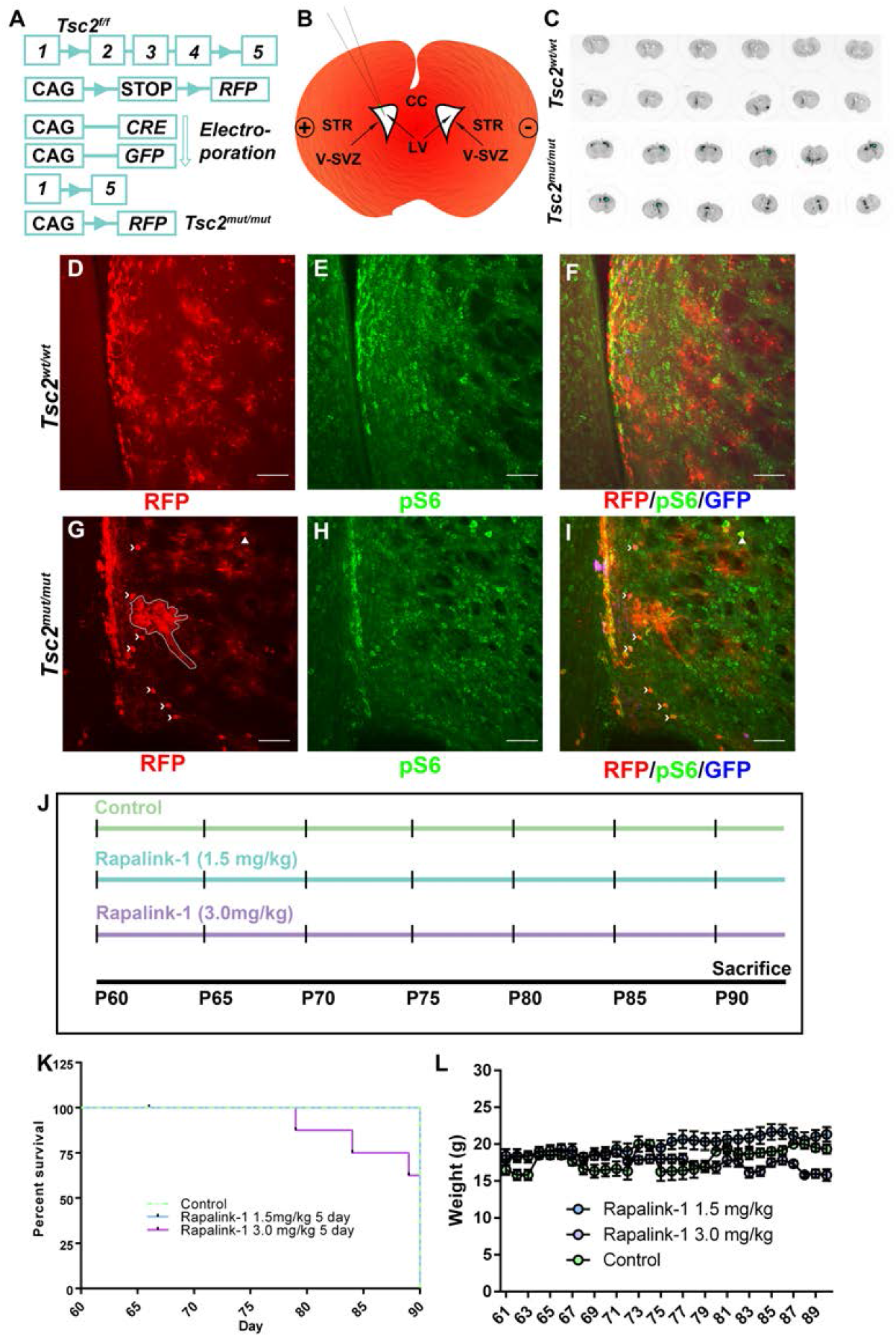
Rapalink-1 Treatment in a TSC Mouse Model. A) Schematic diagram of conditional *Tsc2* and inducible *RFP* genes. CRE recombinase causes *Tsc2* exons 2-4 to recombine, and RFP to be expressed. B) Schematic diagram of CAG-CRE and CAG-GFP (Green) plasmid electroporation. Plasmid is injected intraventricularly into P0-1 mice. Electrodes placed on the head of the pup subsequently cause plasmids to be taken up by NSCs lining the lateral ventricles when an electrical field is applied. C) Macroscopic images of coronal sections of P60 mice following electroporation. Black indicates RFP expression whereas blue indicates RFP saturation. D-F) Images of 20× *Tsc2*^*wt/wt*^ coronal section demonstrating induction of RFP (red, D), pS6 staining (green, E), and GFP expression (blue, F). G-I) Images of 20× *Tsc2*^*mut/mut*^ coronal section demonstrating induction of RFP (red, G), pS6 staining (green, H), and GFP expression (blue, I). Chevrons indicate macroscopic neurons, and arrowhead indicates a giant cell. J) Dose schedule K) Kaplan-Meier survival curve of control and Rapalink-1 treated mice. L) Weights of control and Rapalink-1 treated mice. Data is represented as mean ± SEM. Scale bar = 75 μm.

### Rapalink-1 inhibits mTORC1 activity

mTORC1 activity and substrate phosphorylation in the brain is cell type dependent^29–31^. We noted that pS6 levels were high within RFP positive *Tsc2* null cells having neuron-like morphologies (**Figure 2**)^27^. pS6 was analyzed in Control, Rapalink-1 (1.5 mg/kg), or Rapalink-1 (3.0 mg/kg) *Tsc2* mutant neurons. We found that Rapalink-1 decreased pS6 (DMSO Mean=1.066±0.029 n=857 vs. Rapalink-1 1.5 mg/kg Mean=0.9127±0.065 n=218 vs. Rapalink-1 3.0 mg/kg Mean=0.835±0.029 n=452, p<0.0001) (**Figure 2G**).

**Figure 2.**
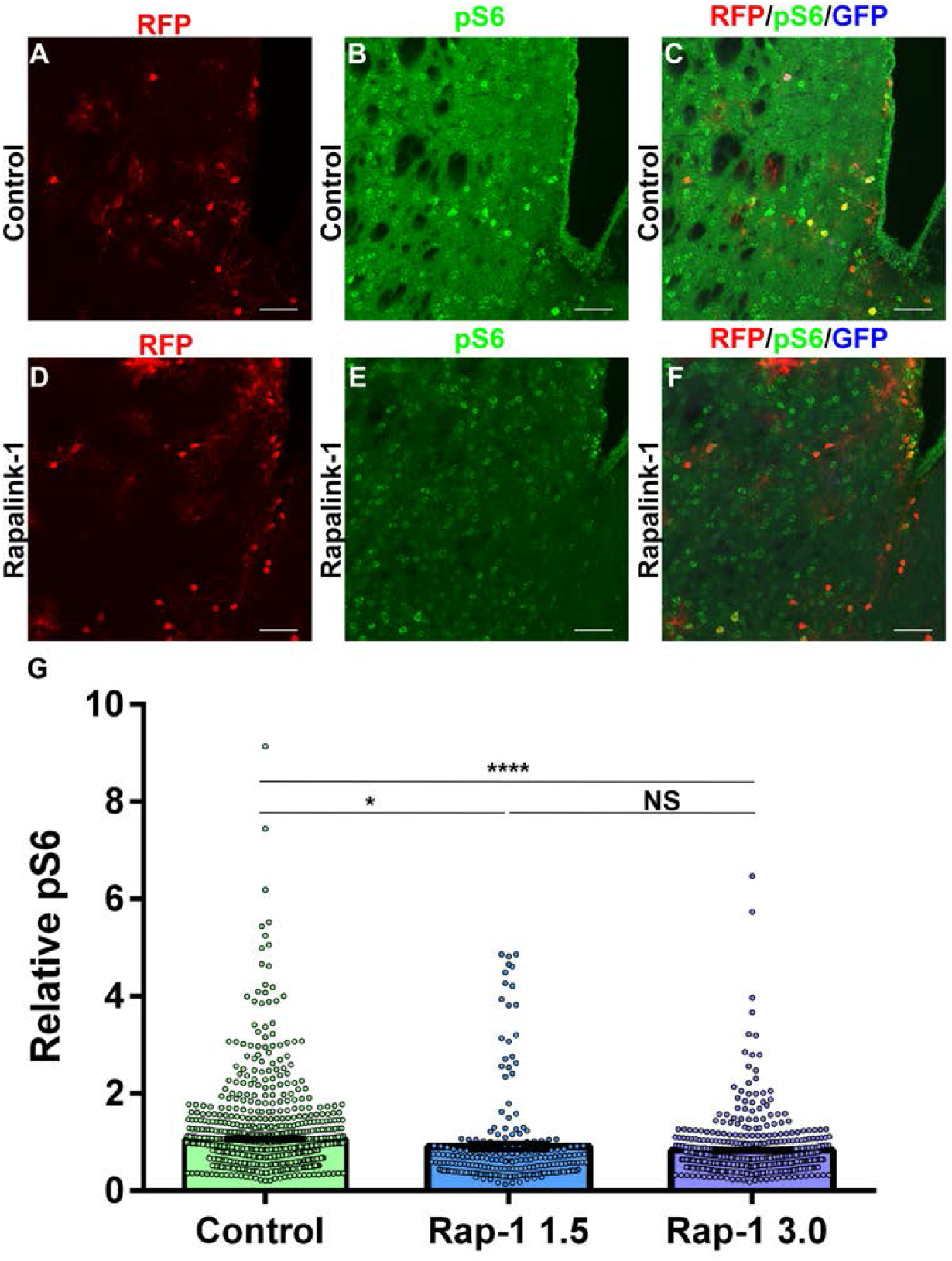
Rapalink-1 Inhibits mTORC1 Activity. A-C) 20× images of RFP (red, A), GFP (blue, B), and pS6 (green, C) in a coronal section of a *Tsc2*^*mut/mut*^ P90 control brain. D-F) 20× images of RFP (red, D), GFP (blue, E), and pS6 (green, F) in a coronal section of a brain from a *Tsc2*^*mut/mut*^ P90 Rapalink-1 (3 mg/kg) treated mouse. G) Quantification of pS6 in RFP positive cells. *=p<0.05, ****=p<0.0001. Data is represented as mean ± SEM. Scale bar = 75 μm.

We confirmed that mTORC1 signaling was reduced by using acute daily Rapalink-1 treatment for five days. Organs were harvested following 6 mg/kg Rapalink-1. Rapalink-1 reduced mTORC1 signaling as detected by p4EBP staining and by immunoblotting for pS6 (Control = 1.000 ± 0.04189, N=3 vs Rapalink-1 = 0.7707 ± 0.02518, N=3; P=0.0094) (**Supplemental Figure 1**). These results confirm that Rapalink-1 can indeed reduce mTOR signaling *in vivo*.

### Rapalink-1 reduces Tsc2 mutant neuron cell size

TOR regulates olfactory bulb granule cell soma growth^32^. Loss of mTOR and inhibition of mTORC1 with rapamycin decreases granule cell soma size^32^. Conversely, loss of *Tsc2* increases granule cell and striatal neuron soma size^27,31^. We wondered to what extent soma size might be decreased in *Tsc2* mutant neurons following Rapalink-1 treatment. Rapalink-1 low dose (1.5 mg/kg) and Rapalink-1 high dose (3.0 mg/kg decreased) the average soma size of mutant *Tsc2* neurons (Control Mean=1.00±0.016 vs. Rapalink-1 1.5 mg/kg Mean=0.7463±0.01 vs. Rapalink-1 3.0 mg/kg Mean=0.689±0.015, p<0.0001) (**Figure 3A-H, K**). We documented a population of cells appearing as giant cells having high mTORC1 activity even after Rapalink-1 treatment (**Figure 3I, J**). However, the relative proportion of cells that were classified as giant cells in relation to the total cell number per section, was reduced in the high dose Rapalink-1 condition (Control Mean=0.0451±.0044 vs. Rapalink-1 1.5 mg/kg Mean=0.0415±0.0125 vs. Rapalink-1 3.0 mg/kg Mean=0.0274±0.03623, p<0.01). Taken together, Rapalink-1 appears to decrease the average size of *Tsc2* mutant neurons.

**Figure 3.**
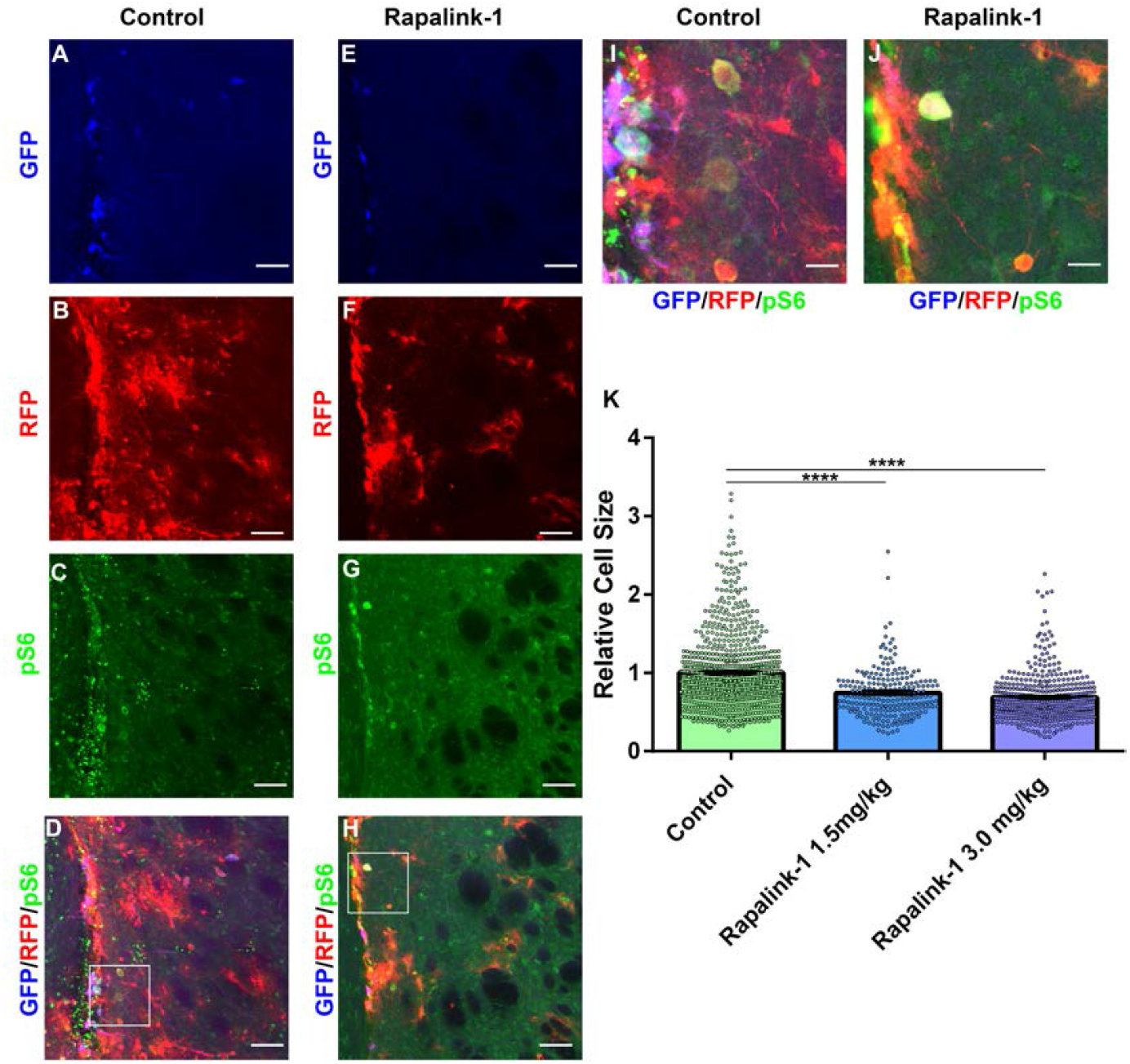
Rapalink-1 Reduces Cell Size. A-D) 20× images of GFP (blue, A), RFP (red, B), pS6 (green, C), and composite (D) in *Tsc2*^*mut/mut*^ P90 control brains. E-H) 20× images of GFP (blue, E), RFP (red, F), pS6 (green, G), and composite (H) in *Tsc2*^*mut/mut*^ P90 Rapalink-1 (3 mg/kg) brains. I, J) Images from D and H magnified 400% showing examples of giant cells. K) Quantification of neuron soma size relative to controls. Data is represented as mean ± SEM. Scale bar = 75 μm. ****=p<0.0001

### Rapalink-1 Treatment Decreases Neuron Dendrite Arbors

We previously demonstrated that loss of *Tsc1, Tsc2*, or increasing Rheb, increased dendrite arbors of olfactory bulb granule cell neurons produced from V-SVZ NSCs^31,33,34^. This is likely because mTOR and mTOR complex components raptor and rictor regulate granule cell dendrite arbors^32^. Neurons present in the striatum of the SEGA model also have more dendrites than wild type or *Tsc2* mutant olfactory bulb granule cell neurons^27^. These mutant neurons are also larger and have greater dendrite complexity in comparison to any control wild-type neurons found in the striatum.

Gross reductions in dendrite arbors caused by Rapalink-1 were noted at low magnifications (**Figure 4A, B**). To confirm and quantify this observation, neurons from control and Rapalink-1 treated mice were traced (**Figure 4C, D**). Rapalink-1 reduced the length and complexity of *Tsc2*^*mut/mut*^ neuron dendrites (control = 713.0 ± 74.53, n=24 vs. Rapalink-1 (3.0 mg/kg) 465.5 ± 33.85, n=50; p=0.008) (**Figure 4E, F**). Thus, long-term Rapalink-1 treatment significantly reduces dendrite arbors in *Tsc2* mutant neurons.

**Figure 4.**
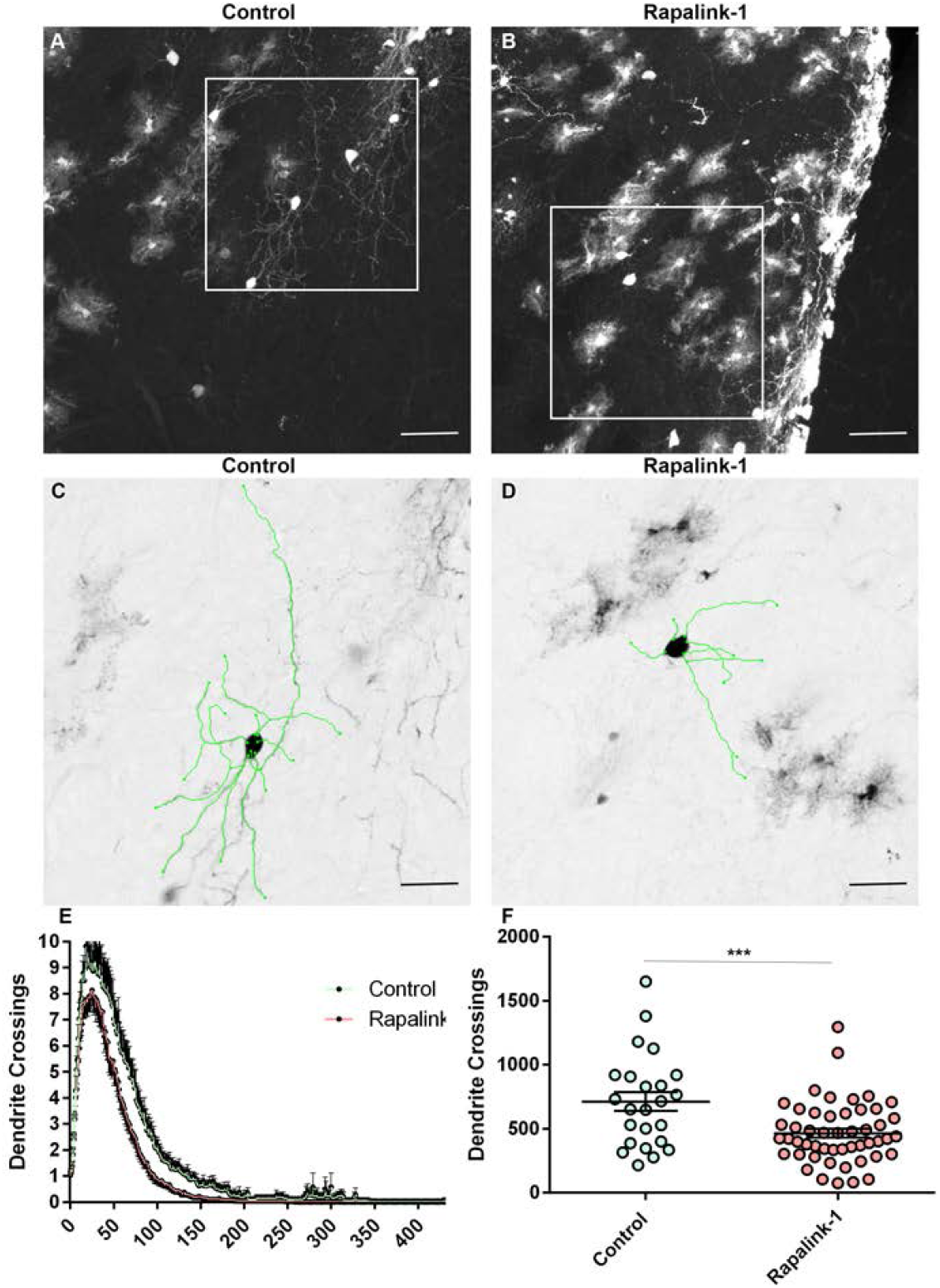
Rapalink-1 Reduces Dendrite Arbors in a TSC Model. A) 20× image of RFP in a *Tsc2*^*mut/mut*^ P90 control brain. B) 20× image of RFP in a *Tsc2*^*mut/mut*^ P90 Rapalink-1 treated brain. C) Zoomed in image with a neuron in square of A subjected to tracing (green highlight). D) Zoomed in image with a neuron in square of B subjected to tracing (green highlight). E) Sholl Analysis demonstrating the effect of Rapalink-1 on dendrite arbors. F) The total number of dendrite crossings in each neuron of control and Rapalink-1 treated mice. Data is represented as mean ± SEM. Scale bar = 75 μm (A, B) or 37.5 μm (C, D). ***=p<0.001

### Efficacy of Rapalink-1 for the Treatment of TSC Hamartomas

The final determinant of the efficacy of Rapalink-1 was the average lesion size at P90. Striatal hamartomas were measured by tracing lesions similar to what we previously described^27^. We noted that size continues to increase notably at P90 with some hamartomas appearing aggressive and invading different regions reaching 10 times the size of those measured at P60 (**Figure 1, 5**). In comparison, Rapalink-1 significantly reduced the hamartoma size in relation to control conditions (control = 40096 ± 8461, n=38 vs. Rapalink-1 3.0 mg/kg 19636 ± 3266, n=33, p=0.0363) (**Figure 5I**).

**Figure 5.**
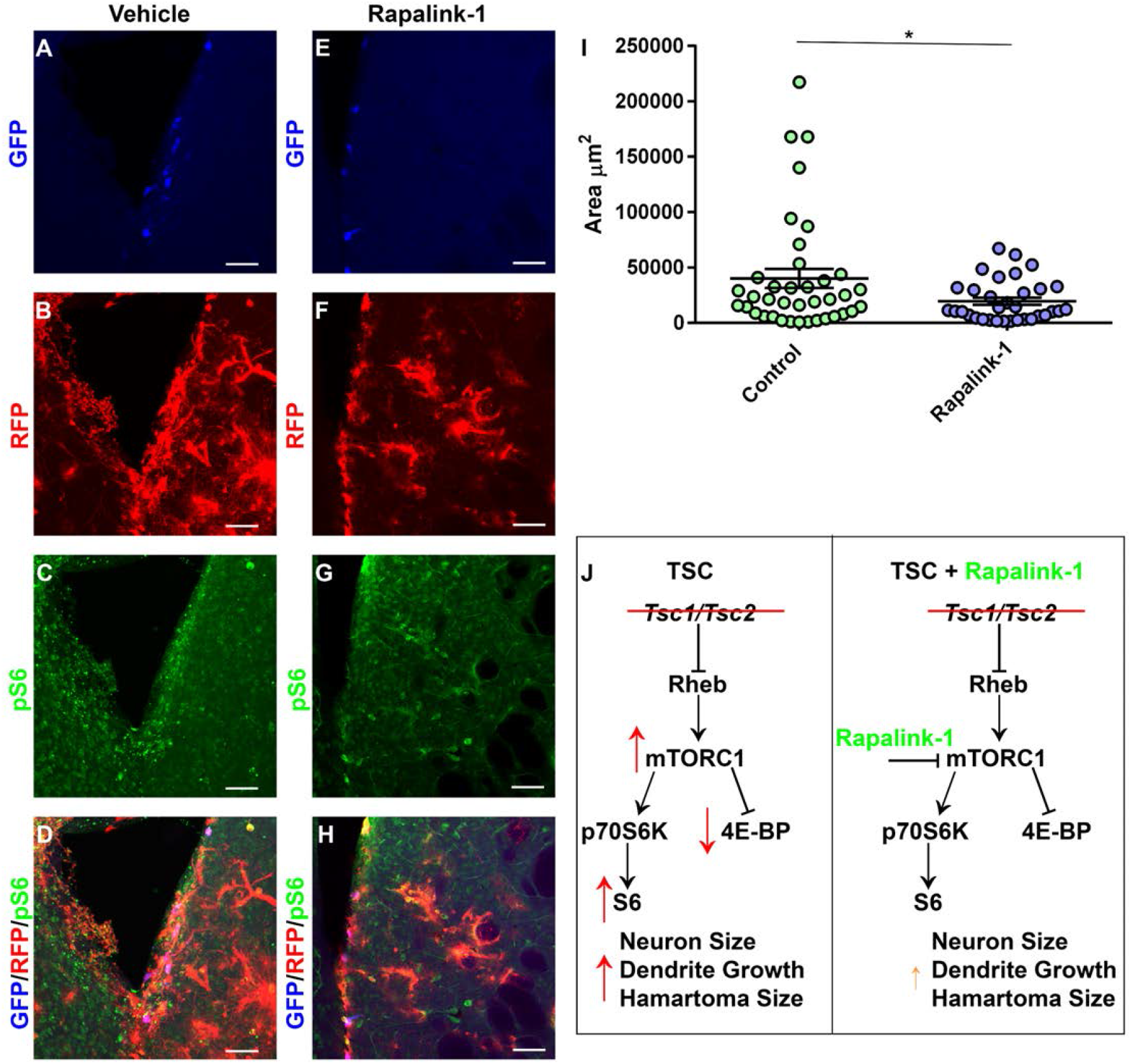
Rapalink-1 Reduces Hamartoma Size. A-D) 20× images of GFP (blue), pS6 (green), and RFP (red) in *Tsc2*^*mut/mut*^ P90 control brains. E-H) 20× images of GFP (blue), pS6 (green), and RFP (red) in *Tsc2*^*mut/mut*^ P90 Rapalink-1 treated brains. I) Quantification of lesion area. J) Schematic diagram of molecular pathways regulated by *Tsc1/Tsc2* in TSC model in control and Rapalink-1 treated brains. Tsc1/Tsc2 turn off mTORC1 pathway under normal conditions. However, in the TSC SEGA model (left), *Tsc1/Tsc2* encoded protein products cannot turn mTORC1 off leading to ectopic/cytomegalic neurons with excessive mTORC1 activity, hypertrophic dendrites, giant cells, and SEGA-like lesions (i.e. striatal hamartomas). Rapalink-1 partially rescued these changes and inhibited the mTORC1 pathway activity, decreased neuron size, and reduced SEGA-like lesions. Data are represented as mean ± SEM. Scale bar = 37.5 μm *=p<0.05

## Discussion

SEGAs are a significant cause of morbidity in TSC patients. Rapalogs are a front-line treatment for SEGAs^35–38^. Here we demonstrate that Rapalink-1 is useful for reversing neural phenotypes in a TSC mouse model of SEGA-like hamartomas (**Figure 5J**). Although, ∼57% of TSC SEGAs are reduced by 50% within two years, continued treatment does not further reduce SEGA volume^18,19,22^. Protracted use of Rapalogs is also necessary because stopping treatment is associated with SEGA recurrence. Thus considering the overall moderate efficacy of Rapalogs and in consideration of the side effects, especially for children, warrants that new medications are deployed^23^. There is an urgent need to test new therapies in model systems. Rapalink-1 is an excellent candidate because it fulfills the need to completely inhibit mTORC1 phosphorylation of rapamycin-resistant substrates and simultaneously inhibits mTORC2^26^. To fulfill the goal of examining additional TSC therapies, we tested Rapalink-1 in a mouse model of TSC SEGA-like hamartomas.

The mechanisms that account for SEGA growth are unclear in patients and this may be related to the fact that we have little understanding of the cellular composition, regional differences in SEGAs, and whether molecular mechanisms involved in the different regions and timing of their appearance may differ. The fact that the composition of SEGAs has been debated for nearly a century is evidence that experiments must carefully examine the effect drugs have not just on SEGA size, but also on the different cell types within SEGAs^39,40^. SEGAs also appear to change over time. Imaging and RNA sequencing experiments suggest a stepwise change in composition with SEGAs eventually containing neurons and cytomegalic giant cells^41–47^. Gliotic scarring, calcification, and immune cell invasion also occur^47^. This is similar to events in TSC patient cortical tubers^48^. Although SEGAs do not frequently directly cause seizures, the presence of mutant neurons and giant cells warrants closer inspection as does determining their contributions to patient presentation.

Loss of *Tsc2* in V-SVZ neuroprogenitors results in the aberrant transition from quiescent to active states and during differentiation^29,30,49,50^. Loss of *Tsc2* prevents the further downregulation of mTORC1 activity in a subset of cells within the striatum leading to growths including nodular hamartomas containing cells with heterogenous morphologies. The hamartomas are characterized by abnormal heterotopic clusters of morphologically heterogenous cells which include neurons both within and outside of the growths. Thus, the TSC SEGA-like hamartoma mouse model allowed us to examine the utility of Rapalink-1 treatment in controlling neural phenotypes and hamartoma size. Rapalink-1 intermittent dosing (1.5 mg/kg and 3.0 mg/kg every five days) was generally tolerated during the study, but preliminary experiments demonstrated acute toxicity during daily treatment (data not shown). As predicted, Rapalink-1 reduced mTORC1 activity. However, it was reduced by only ∼15-23% as measured using pS6 as a readout and pS6 remained apparent throughout the brain. Thus, these doses might incompletely inhibit mTORC1. Indeed, high dose (6 mg/kg) appeared more effective by immunohistochemistry and western blot but was associated with toxicity. Additionally, 1.5 and 3.0 mg/kg Rapalink-1 treated mice were sacrificed 5 days after the last injection of Rapalink-1. Thus, mTORC1 activity could have rebound effects and our data underestimate how effective Rapalink-1 is. Nevertheless, the reduction in pS6 served as an indicator that Rapalink-1 inhibited mTORC1.

We reasoned that the prolonged inhibition of mTORC1 would decrease the well-documented mTORC1-regulated phenomenon, of cell growth^51^. Previous work from our laboratory and our colleagues have demonstrated that neonatal V-SVZ NSC generated granule cells grow when *Tsc1* or *Tsc2* are removed^31,33^. Moreover, ectopic expression of wild-type or mutant Rheb can drive cell growth in granule cells^34,52^. Growth of granule cells is dependent on mTOR since CRE electroporation of conditional mTOR reduces soma size^32^. Moreover, rapamycin treatment reduces granule cell soma size supporting the importance of mTORC1 in this process^32^. As expected, Rapalink-1 at both doses reduced *Tsc2* mutant neuron cell size by ∼25-31%. These results further support that Rapalink-1 is likely inhibiting mTORC1-dependent cellular events.

While the role of striatal neurons in the hamartoma SEGA-like model is unclear, ectopic neurons can affect a wide range of other cell types. Cortical tubers and focal malformations of cortical development appear to undergo analogous changes including astrogliosis and microglia activation^11^. And this is directly linked to neuron hyperexcitability. Thus, a major goal for TSC and related disorder research has been to reduce dendrite growth in the hopes of modulating neuronal activity. mTOR, through mTORC1 and mTORC2 regulates olfactory bulb granule cell dendrite arbors^32^. While cytoskeleton regulation is most often attributed to mTORC2, rapamycin and mTORC1 have been carefully studied in relation to dendrite growth. For example, like cell growth, loss of *Tsc1* and *Tsc2* as well as increased Rheb activity promote dendrite growth^31,33,34,52^. We found that Rapalink-1 similarly reduced the dendrites of striatal *Tsc2* mutant neurons. Thus, Rapalink-1, could be advantageous compared to rapalogs in that it binds both arms of mTOR signaling^53^.

This is particularly important in that several groups have posited that additional pathways may regulate SEGA growth. For example, one group simultaneously removed *Pten* and *Tsc1* in postnatal V-SVZ NSCs^16^. This resulted in the generation of SEGAs that recapitulated most aspects of those in patients. *Pten/Tsc1* mutant NSCs were subsequently injected subcutaneously and generated tumors. The *Pten*/*Tsc1* mutant NSCs had altered Erk and Akt activity too. Knockdown of the mTORC2 component rictor or combined rapamycin and PI3K-mTOR inhibition reduced tumor growth^16^. Thus, inhibitors that act on both mTORC1 and mTORC2 such as Rapalink-1 have several advantages. In line with these results, we found that Rapalink-1 produced a moderate but significant effect on the average size of SEGAs. The effect of Rapalink-1 was disproportionate to that seen on cell size or dendrites. Whether the effectiveness of Rapalink-1 is due to the effect on neuron activity or on other cell types is unclear and will require additional future experiments.

Taken together, this study provides *in vivo* evidence for the utility of Rapalink-1 to control mTOR, cell size, dendrite hypertrophy, and SEGA-like hamartoma growth which may be of clinical importance.

## Acknowledgements

This work was supported by the United States of America Department of Defense U.S. Army Medical Research Activity Award Congressionally Directed Medical Research Program Tuberous Sclerosis Complex Research Program W81XWH2010447.

## Author Contributions

Conceptualization, D.F., Methodology, S.M., D.F., M.W., M.S., V.R.; Validation, D.F., V.R., M.W., M.S.; Formal Analysis, S.M., M.W., M.S., D.F.; Investigation, D.F., S.M., V.R., M.W., M.S.; Resources, D.F.; Data Curation, D.F., M.W., M.S.; Writing-Original Draft, D.F.; Writing-Reviewing and Editing; D.F., S.M., V.R., M.W., M.S.; Visualization, S.M., D.F.; Supervision, D.F.; Project Administration, D.F.; Funding Acquisition, D.F.

## Figure Legends

**Supplementary Figure 1.** *Acute high-dose Rapalink-1*. A-B) Example of P60 p4EBP1 (green) staining. Note the presence of p4EBP1 in lesion cells (RFP, red) within the striatum. C-D) Example of P90 p4EBP1 (green) staining. Note the presence of p4EBP1 in lesion cells (RFP, red) within the striatum. E-F) Example of ∼P66 p4EBP1 (green) staining following Rapalink-1 treatment (6mg/kg) daily. Note the absence or diminished p4EBP1 (green) in lesion cells (RFP, red) within the striatum. G) Western blot of pS6, total S6 (short exposure), total S6 (long exposure of same length), and GAPDH from hearts of mice treated with DMSO or high dose Rapalink-1. All western blots have not been spliced. H) Quantification of western blot data. Note that while pS6/S6 was decreased, both the phosphorylated form and total form of S6 appeared lower. Data is represented as mean ± SEM. Scale bar = 75 μm. *=p<0.05

## Methods

### EXPERIMENTAL MODEL AND SUBJECT DETAILS

#### Animals

All experiments were approved by the Clemson University Institutional Animal Care and Use Committee and the Animal Care and Use Review Office (ACURO), a component of the USAMRDC Office of Research Protections (ORP) within the Department of Defense (DoD). All methods were carried out in accordance with relevant guidelines and regulations. All methods are reported in accordance with ARRIVE guidelines. Red fluorescent protein (RFP^+/-^,^+/+^) (B6.Cg-Gt(ROSA)26Sortm9(CAG-tdTomato)^Hze/J^) (Strain #007909, RRID:IMSR_JAX:007909), and *Tsc2*^tm1.1Mjg/J^ (Strain #027458, RRID:IMSR_JAX:027458) were acquired from Jackson Laboratories^54^. Sentinel mice were free of pathogens throughout the study. Samples/subjects were allocated randomly to experimental group. Experimental manipulations were performed on mouse pups that were not involved in previous procedures and sacrificed accordingly. Ages and both sexes were used as indicated in figures. Mice were housed under standard pathogen-free conditions in cages on racks within isolated cubicles with a 12-h light/dark cycle and fed *ad libitum*. Mice were injected intraperitoneally using the indicated doses and schedule. Drugs were prepared as previously described in DMSO, PEG-300, and PBS^53^. Injections were performed in biological safety cabinets. Mice were weighed 5-7 days each week and weights recorded. Animals were euthanized by intraperitoneal injection of euthasol or CO_2_ inhalation followed by swift decapitation.

#### Electroporation

B6.Cg-Gt(ROSA)26Sortm9(CAG-tdTomato)^Hze/J^) x *Tsc2*^tm1.1Mjg/J^ mouse pups were electroporated as previously described^55,56^. Mice were injected with equal concentrations and volumes of DNA plasmids diluted in phosphate buffered saline (PBS) with 0.1% fast green. CAG-CRE (Plasmid #13775, Addgene) and CAG-GFP (Plasmid #11150, Addgene) plasmids were used^57,58^. A borosilicate glass micropipette generated from pulled capillary tubes was loaded with DNA and injected into the lateral ventricles. Square pulse generation was performed using a pulse generator (ECM830; BTX) and tweezer electrodes (model 520; BTX) with five, 100-volt square pulses of 50 ms duration with 950-ms intervals.

#### Polymerase Chain Reaction (PCR)

Toe or tail snips were subject to modified hotshot DNA extraction and genotyped by PCR using Invitrogen™ Platinum™ Taq polymerase, mixed with nuclease-free water, magnesium free PCR buffer, MgCl_2_, dNTP mix with primers and DNA according to the manufacturers’ protocol (Invitrogen). Primers for *Tsc2* were 5’-ACAATGGGAGGCACATTACC-3’ and 5-AAGCAGCAGGTCTGCAGTG-3’ and for Tomato (RFP) 5’-AAGGGAGCTGCAGTGGAGTA-3’ and 5’-CCGAAAATCTGTGGGAAGTC-3’ and 5’-GGCATTAAAGCAGCGTATCC-3’ and 5’-CTGTTCCTGTACGGCATGG-3’. Amplicons were loaded onto agarose gels with 1X Blue Juice and ran at 100 V for 20–30 mins and visualized on a BioRad Chemidoc MP.

#### Immunohistochemistry

Brains were removed in room temperature PBS, transferred to 4% paraformaldehyde in PBS, and incubated overnight at 4°C. Brains were rinsed in PBS and mounted in 3% agarose. A Leica VTS 1000 vibratome was used to section brains coronally. Sections were blocked in 0.1% Triton X-100, 0.1% Tween-20 and 2% BSA in PBS for 1 hr at room temperature. Sections were washed in 0.1% Tween-20 in PBS. Sections were incubated in primary antibody, anti-pS6 (1:500; Cell Signaling Technology; Ser 240/244, 61H9, #4838), in 0.1% Tween-20 and 2% BSA in PBS overnight at 4°C. Sections were subjected to three additional washes in PBS containing 0.1% Tween-20. Sections were incubated with the appropriate secondary antibody (Alexa Fluor series; 1:500; Invitrogen) in 0.1% Tween-20 and 2% BSA in PBS overnight at 4°C. Sections were mounted in ProLong Antifade Mountant (Thermo Fisher Scientific). Images were acquired on a spectral confocal microscope (Leica SPE) with a ×20 dry objective (N.A. 0.75). Low-magnification images were acquired with ×10 dry or a ×5 dry (N.A. 0.15) objective.

#### Image Analysis

Images (×20) of RFP positive cells were uploaded to FIJI (ImageJ 1.5 g) and analyzed as described elsewhere^27^. The freehand selection tool was used to trace electroporated and non-electroporated cell somas in the same Z section and mean gray values for pS6 were quantified. Ratios of electroporated and non-electroporated cells were compared for RFP positive cells in *Tsc2*^*wt/wt*^ and *Tsc2*^*mut/mut*^ conditions to account for immunohistochemical variation. Soma size was simultaneously recorded for traced cells. The RFP positive hamartoma perimeter was outlined in each Z section by hand by scrolling through individual Z sections and hamartomas were subsequently traced in (×20) images. The freehand selection tool was subsequently used to trace the margins of the complete hamartoma within the striatum. Images (×20) were used to measure dendrite morphology. Dendrites were traced using the simple neurite tracer plug-in. Sholl analysis was performed at 1 μm intervals to quantify dendrite arborization using the Sholl plug-in. The total number of dendritic crossings was calculated by taking the sum of crossings at all intervals for each traced neuron and averaging the total number of crossings per neuron in each condition.

#### Western Blot

Tissue (0.1 gram) was harvested and finely minced in 2% SDS, Protease Inhibitor Cocktail (Pierce) and Phosphatase Inhibitor Cocktail, in RIPA buffer. Samples were briefly sonicated at maximum settings (100 Amplitude, QSonica) for 10 seconds three times with 30 second resting intervals. Lysate was transferred on ice to a fresh reaction tube and centrifuged at 15,000 rpm for 15 min in a tabletop Eppendorf 5415 centrifuge at 4°C. Protein concentration was quantified using the Pierce MicroBCA assay. Equal protein amounts were brought up to equal volumes with lysis buffer as described above and Laemmli buffer and heated to 95°C for 5 minutes. Proteins were resolved by 10% polyacrylamide precast mini-Protean gels (BioRad) and transferred to polyvinylidene difluoride (PVDF) membranes. PVDF membranes were rinsed in Tris-buffered saline (TBS-T, 0.1% Tween 20) for 5 min at room temperature and blocked in 5% blotting grade block (BioRad) in TBS-T for 1 h at room temperature. Membranes were incubated for 1 h at room temperature or overnight at 4°C with the following antibodies from Cell Signaling Technology at a 1:1,000 dilution: phospho-RPS6 (D68F8, Cat# 5364) and RPS6 (5G10, Cat# 2217). Membranes were rinsed three times each for 10 min in TBS-T. Membranes were incubated for 1 h at room temperature with donkey or goat anti-rabbit antibodies in blocking buffer. Membranes were washed for 15 minutes in TBS-T and visualized using a Bio-Rad Chemidoc MP imaging system using enhanced chemiluminescence reagent (Pierce). PVDF membranes were stripped for at room temperature using Restore Western Blot Stripping Buffer according to manufacturer’s recommendations (Cat# 21059, Thermo Fisher Scientific).

### QUANTIFICATION AND STATISTICAL ANALYSIS

Measurements were graphed and statistical analysis was performed with GraphPad Prism software (Version 8.2.0, GraphPad Software Inc.). Statistical significance was determined using One-way analysis of variance (ANOVA) with Tukey’s multiple comparisons test or Student’s T-test. N (number of mice) and n (number of cells or hamartomas) are listed where applicable. Error bars are reported as the standard error mean.

### KEY RESOURCES

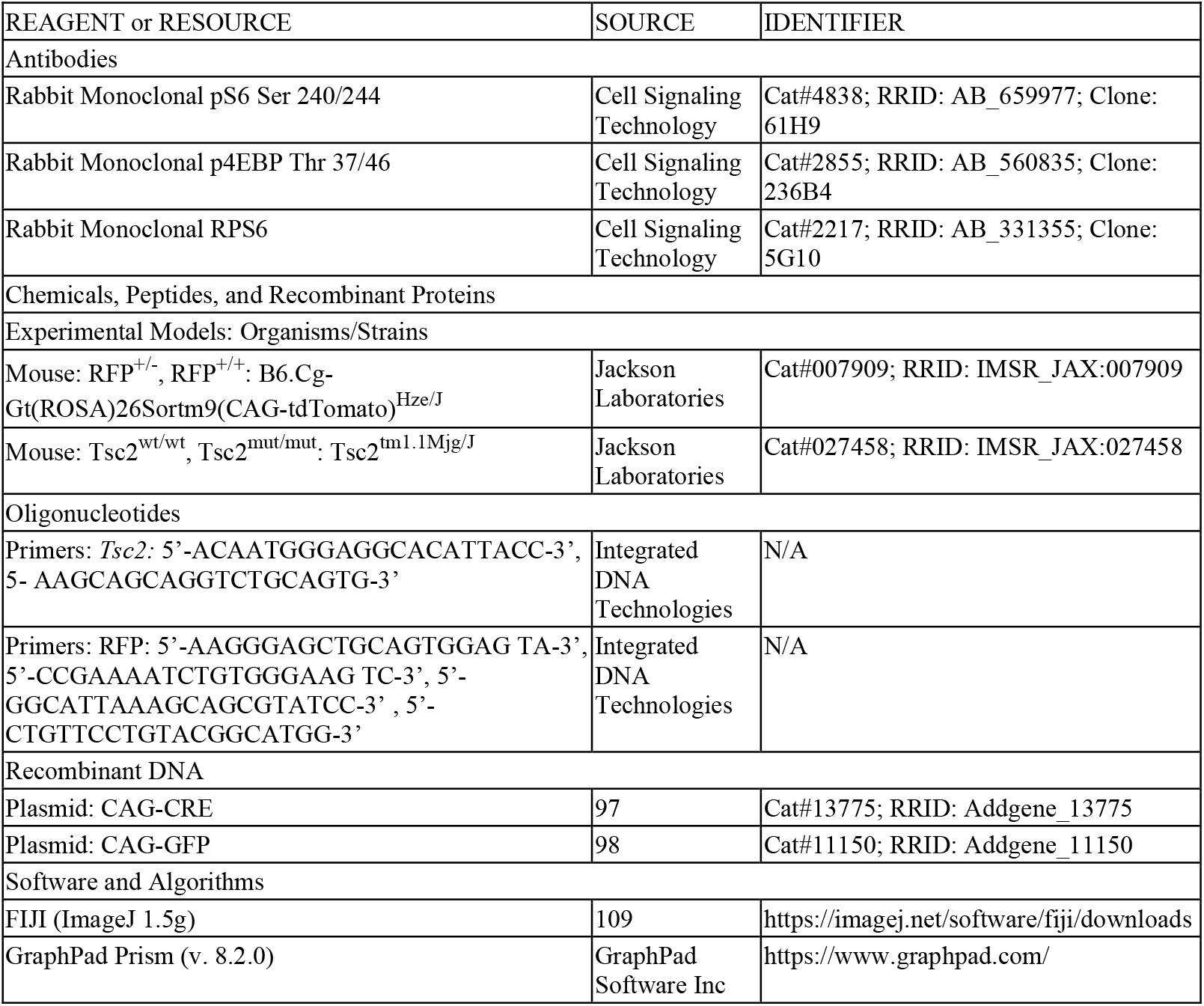

## DATA AVAILABILITY

The datasets generated and/or analysed during the current study are available in the Mendeley Data and NeuroMorpho repository. Western blot data is deposited as Feliciano, David (2025), “A Bitopic mTORC Inhibitor Reverses Phenotypes in a Tuberous Sclerosis Complex Model Westerns”, Mendeley Data, V1, doi: 10.17632/hjt9m462d3.1 (https://data.mendeley.com/datasets/hjt9m462d3/1) to Mendeley Data. Neuron traces are available at NeuroMorpho.org under “A Bitopic mTORC Inhibitor Reverses Phenotypes in a Tuberous Sclerosis Complex Model” or navigating to neuromorpho.org/dableFiles/mukherjee_feliciano/Supplementary/Mukherjee_Feliciano.zip.

